# Lyophilization Maintains the Storage Stability and Bioactivity of Mesenchymal Stem Cell–Derived Extracellular Vesicles

**DOI:** 10.64898/2026.06.10.731407

**Authors:** Young-Ju Lim, Wook Tae Park, Joo-Hee Choi, Gun Woo Lee, Min-Jung Ma, Wansuk Son, Seunghyun Kang, Min-Soo Seo, Sangbum Park

## Abstract

**Background:** Extracellular vesicles (EVs) derived from mesenchymal stem cells (MSCs) are emerging therapeutic candidates for bone regeneration, but their long-term stability remains a barrier to clinical translation. This study evaluated whether lyophilization supports the stable storage of extracellular vesicles derived from human epidural fat MSCs (hEF-MSCs) cell line.

**Research Design and Methods:** EVs were isolated, lyophilized with 8.5 % sucrose and HEPES(4-(2-Hydroxyethyl)piperazine-1-ethanesulfonic acid, N-(2-Hydroxyethyl)piperazine-N′-(2-ethanesulfonic acid)), and stored for 3 months at −80, −20, 4, and room temperature (20). After reconstitution in phosphate-buffered saline (PBS), structural characteristics were assessed using transmission electron microscopy, nanoparticle tracking analysis, and flow cytometry. Functional activity was evaluated using MC3T3-E1 osteogenic differentiation assays. The main outcome measures included morphology, particle counts, marker expression, cytotoxicity, and sequencing profiles.

**Results:** Lyophilization maintained EV morphology, structural integrity, and particle distribution across all storage temperatures. Marker expression remained comparable among the conditions. Reconstituted EVs promoted osteogenic differentiation of MC3T3-E1 cells without evidence of cytotoxicity. Sequencing profiles revealed no significant differences among storage conditions.

**Conclusions:** Lyophilized EVs from hEF-MSCs cell lines exhibited stable structural and functional properties across a range of storage temperatures, supporting their suitability for further development in bone regeneration applications.

## 1. Introduction

Nanomedicine has rapidly advanced as a central technology in modern biomedicine, particularly through the development of lipid-based delivery platforms such as liposomes. The first clinically approved nanoscale anticancer formulations, introduced in the 1990s, were liposomal drugs encapsulating doxorubicin (Barenholz, 2012; van der Meel et al., 2013). More recently, several effective COVID-19 vaccines have employed mRNA encapsulated within lipid nanoparticles (Shi et al., 2024; Swetha et al., 2023). These advanced nanocarriers are artificially engineered vesicles composed of synthetic lipid bilayers that frequently incorporate cationic lipids to enhance interactions with cellular membranes and are often coated with polyethylene glycol (PEG) to prolong circulation by limiting immune detection (Haghighi et al., 2024; Sun and Lu, 2023). Despite their utility, liposomal systems exhibit notable limitations. Cationic lipids may contribute to cytotoxicity, and repeated exposure to PEG-modified surfaces can trigger anti-PEG antibody responses (Alton et al., 2014; Lv et al., 2006; Shi et al., 2022; Zhang et al., 2016). Moreover, their *in vivo* biodistribution is difficult to regulate without the use of specific targeting ligands, which substantially increases manufacturing complexity and cost. These challenges have prompted interest in naturally derived extracellular vesicles (EVs) as promising alternatives for next-generation drug delivery platforms.

EVs have emerged as key mediators of intercellular communication, transporting a diverse array of bioactive molecules, including proteins, lipids, and nucleic acids (Kalluri and LeBleu, 2020; Yáñez-Mó et al., 2015; Zhang et al., 2019). Increasing evidence shows their therapeutic potential across multiple biomedical fields, from oncology to regenerative medicine. EVs are particularly attractive as alternatives to cell-based therapies because they replicate many of the beneficial paracrine effects of stem cells while avoiding risks associated with cellular transplantation, such as tumorigenicity and immune rejection (Schlesinger et al., 2017; Weisel et al., 2020). Among their various applications, EVs have shown significant promise in promoting bone regeneration and osteogenic differentiation, addressing critical clinical challenges such as osteoporosis and delayed fracture healing (Biswas et al., 2025; Zhang et al., 2024).

Despite their therapeutic potential, a major barrier to the translational application of EVs is maintaining their structural integrity and biological activity during storage. Conventionally, EVs are stored at −80°C or 4°C, which preserves functionality but is impractical for large-scale clinical deployment due to high costs, stringent cold-chain requirements, and logistical challenges in global distribution. Lyophilization has emerged as a practical approach to address these limitations, enabling long-term storage at higher temperatures while simplifying transport and handling. Previous studies have indicated that lyophilization, particularly when combined with stabilizers such as various oligosaccharides, such as mannitol, trehalose, and sucrose, can maintain vesicle morphology and biological function (Driscoll et al., 2022; Kang et al., 2024). However, inconsistent results and limited evaluations across biological models have raised uncertainties regarding the generalizability of this approach. Among these stabilizers, sucrose is widely used in the lyophilization of lipid nanostructures due to its natural, nontoxic properties (Oh and Gomez, 2018). Furthermore, sucrose is approved by the U.S. Food and Drug Administration (FDA) and conforms to good manufacturing practices (GMP) for human use (Guimarães et al., 2019; Morenweiser, 2005; Rao et al., 2020).

Addressing these challenges requires rigorous evaluation of whether lyophilization preserves EV functionality across clinically relevant storage conditions and durations. In this study, we aimed to determine the stability and osteogenic bioactivity of lyophilized EVs stored at multiple temperatures for up to 3 months. Using MC3T3-E1 pre-osteoblast cells as a model system, we compared the ability of lyophilized EVs to promote osteogenic differentiation with that of freshly isolated counterparts. This systematic study provides critical insights for optimizing EV preservation strategies and advancing EV-based therapeutics toward scalable, clinically applicable solutions.

## 2. Materials and Methods

### 2.1 Reagents

Minimum Essential Medium Eagle Alpha (α-MEM), Dulbecco’s Modified Eagle Medium (DMEM), phosphate-buffered saline (PBS), penicillin–streptomycin (P/S), and 0.25% trypsin–ethylenediaminetetraacetic acid were obtained from Gibco-BRL. Fetal bovine serum (FBS) was purchased from MP Biomedicals. EmeraldAmp GT PCR Master Mix was procured from TaKaRa, and AmpiGene qPCR Green Mix Hi-ROX was obtained from Enzo.

### 2.2 Cell cultures

This study was approved by the Institutional Review Board of Yeungnam University Medical Center (IRB No. 2017-07-032). The human epidural adipose tissue-derived mesenchymal stem cell (hEpi-AD-MSC) line was established from human epidural fat tissue at Yeungnam University Medical Center and was previously immortalized as described by Lee et al. (2025). Human dermal fibroblasts (HDFs) were purchased from the American Type Culture Collection (ATCC, Manassas, VA, USA). HDFs and hEpi-AD-MSCs were cultured in low-glucose Dulbecco’s Modified Eagle Medium (DMEM; Gibco, Carlsbad, CA, USA) supplemented with 10% fetal bovine serum (FBS; Gibco, Carlsbad, CA, USA), 1% penicillin–streptomycin (P/S; Gibco), and 25 μg/mL Plasmocin™ (InvivoGen, San Diego, CA, USA). Cells were maintained at 37°C in a humidified atmosphere containing 5% CO, and the culture medium was replaced every 2–3 days. MC3T3-E1 cells were purchased from the American Type Culture Collection (ATCC, Manassas, VA, USA) and cultured in α-minimum essential medium (α-MEM; Gibco, Carlsbad, CA, USA) containing 1.0 g/L glucose and supplemented with 10% fetal bovine serum (FBS; Gibco) and 1% penicillin–streptomycin (P/S; Gibco). Cells were maintained at 37°C in a humidified atmosphere containing 5% CO, and the culture medium was replaced every 2–3 days.

### 2.3 Isolation of EVs

EVs were isolated from the conditioned medium of hEF-MSCs cultures. To prevent contamination from bovine-derived vesicles, cells were maintained in a medium supplemented with 10% exosome-depleted FBS (Gibco) under standard conditions (37°C, 5% COC). Conditioned medium was collected every 48 h, and cellular debris was removed by centrifugation at 300 × *g* for 10 min. The clarified supernatant was then concentrated using a tangential flow filtration system (Pall Corporation, Port Washington, NY, USA) operated at a flow rate of 120 rpm.

### 2.4 Characterization of EVs

Stem cell marker expression was assessed using flow cytometry (Gallios, Beckman Coulter, Brea, CA, USA) with the following antibodies: CD73-PE (BioLegend, San Diego, CA, USA, 344004), CD90-PE (BioLegend, 555596), CD105-PE (Bio-Rad, Hercules, CA, USA, MCA1557), CD34-FITC (BioLegend, 343504), and CD45-FITC (BioLegend, 555482). Exosomal surface marker expression was analyzed using bead-based flow cytometry (Gallios) with antibodies against CD9 (BioLegend, 358259), CD63 (BioLegend, 353003), and CD81 (BioLegend, 349505). To enable surface marker analysis, EVs were cultured for 24 h with 4% aldehyde/sulfate latex beads (Thermo Fisher Scientific, Rockford, IL, USA). EV morphology was examined using transmission electron microscopy (TEM). EVs were adsorbed onto formvar carbon-coated copper grids (Ted Pella Inc., Redding, CA, USA), fixed with 2% paraformaldehyde for 10 min, and air-dried. Particle size distribution and concentration were measured using nanoparticle tracking analysis (NTA) (Nanosight NS300, Malvern Panalytical, Worcestershire, UK) according to the manufacturer’s instructions.

### 2.5 MTT assay

Cell viability was assessed using the MTT (3-[4,5-dimethylthiazol-2-yl]-2,5-diphenyltetrazolium bromide) assay. MC3T3-E1 cells were seeded in 48-well plates and incubated overnight before treatment with EVs for 1–2 days. The medium was then replaced with fresh medium containing 0, 0.3 × 10^8^, 0.5 × 10^8^, 0.8 × 10^8^, 1 × 10^8^, 5 × 10^8^, and 5 × 10^9^ EVs, and cells were incubated for an additional 1–2 days. Subsequently, 0.5 μg/mL MTT (Sigma-Aldrich) was added to each well, and cells were incubated for 1 h at 37°C in a 5% CO_2_ atmosphere. After incubation, 400 μL of dimethyl sulfoxide was added, and absorbance was measured at 570 nm using an Infinite 200 PRO instrument (Tecan). Relative cell viability was calculated at the conclusion of the experiment.

### 2.6 Reverse-transcription PCR

Total RNA was extracted from cells using TRI-solution (Bioscience Technology). RNA concentration was measured using a multiplate reader according to the manufacturer’s instructions. Reverse transcription-PCR (RT-PCR) was performed using 2 µg of total RNA to assess mRNA expression levels. Primer sequences used for RT-PCR are listed in Table 1. Relative gene expression was normalized to the internal controls β-actin and GAPDH. MicroRNA (miRNA) was isolated using the miRNeasy Cell Mini Kit (Qiagen) according to the manufacturer’s instructions. Ten nanograms of RNA were reverse-transcribed using the TaqMan miRNA Reverse Transcription Kit (Thermo Fisher Scientific). Quantitative PCR was performed using the TaqMan miRNA Assay primers (Thermo Fisher Scientific) under the following conditions: 40 cycles of denaturation at 95°C for 1 s, followed by annealing/amplification at 60°C for 20 s.

### 2.7 Western blot analysis

Total protein was extracted using an EzRIPA lysis kit (ATTO Technology, Tokyo, Japan) and centrifuged at 13,000 rpm for 10 min at 4°C. Protein concentrations were determined using a BCA protein assay. Total protein samples were separated using 10% sodium dodecyl sulfate–polyacrylamide gel electrophoresis and transferred to a polyvinylidene fluoride membrane (Roche, Mannheim, Germany). Membranes were washed with Tris-buffered saline containing Tween 20 (TBST), and blocked with 5% skim milk for 2 h. After blocking, the membranes were incubated overnight at 4°C with primary antibodies. Subsequently, the membranes were incubated with secondary antibodies for 2 h at 4°C, and TBST was used for all washes. Primary antibodies against Runx2 and β-actin were obtained from Santa Cruz Biotechnology (CA, USA). Antibodies against Dlx5, CD9, CD81, Flotillin-2, tumor susceptibility gene 101, Calnexin, GAPDH, AMPK, and Smad proteins were obtained from Abcam (Cambridge, MA, USA). Protein signals were detected using an enhanced chemiluminescence reagent (Advansta) according to the manufacturer’s protocol. Densitometric analysis was performed using a FUSION Solo analyzer system (Vilber Lourmat, Eberhardzell, Germany).

### 2.8 MSC differentiation assays

The hEF-MSCs were seeded in 24-well culture plates. For osteogenic differentiation, MSCs were cultured using the StemPro™ Osteogenesis Differentiation Kit (Gibco, Thermo Fisher Scientific, Waltham, MA, USA) for 20 days. After differentiation, cells were fixed with 4% formaldehyde (Sigma-Aldrich) for 5 min at room temperature(20°C) and stained with 2% Alizarin Red solution (ARS; Chemicon, Merck Millipore, Billerica, MA, USA) for 15 min. For adipogenic differentiation, MSCs were cultured with the StemPro™ Adipogenesis Differentiation Kit (Gibco, Thermo Fisher Scientific) for 21 days, fixed with 4% formaldehyde for 10 min, and stained with 0.5% Oil Red O solution (Sigma-Aldrich) for 15 min. For chondrogenic differentiation, MSCs were pelleted by centrifugation at 200 ×*g* for 5 min in 1.5 mL tubes and cultured using the StemPro™ Chondrogenesis Differentiation Kit (Gibco, Thermo Fisher Scientific) for 21 days. Chondrocyte spheroids were fixed in 4% formaldehyde for 5 min at room temperature(20°C) and stained with Alcian Blue 8GX (Sigma-Aldrich) for 30 min. MSCs cultured without differentiation supplements served as controls. All differentiation experiments were performed in duplicate and repeated independently three times.

### 2.9 Alkaline phosphatase activity and mineralization

To assess alkaline phosphatase (ALP) activity and mineralization, MC3T3-E1 cells were seeded into 24-well plates and cultured for 24 h in α-MEM supplemented with 10% FBS. Osteogenic medium was prepared by adding AA (ascorbic acid) and β-glycerophosphate alongside EVs. Cells were maintained in osteogenic medium for either 7 days or 3 weeks. For ALP staining, cells were fixed with 4% formaldehyde (Merck KGaA, Darmstadt, Germany) for 5 min, rinsed twice with deionized water, and incubated with BCIP^®^/NBT solution (Sigma-Aldrich) for 20 min at room temperature(20°C). Mineralization was assessed by staining with 2% ARS (Sigma-Aldrich) for 20 min at room temperature(20°C). After staining, cells were thoroughly washed with deionized water.

### 2.10 Lyophilization

EVs were collected and concentrated from conditioned media as described in the preceding sections. Before lyophilization, EVs were suspended in a protective solution containing 8.5% (w/w) sucrose and 20 mM HEPES to preserve their structural integrity and biological activity. Lyophilization was performed following a modified protocol based on Guarro et al. (2022), which has been shown to maintain EV morphology, particle size distribution, and functional properties after reconstitution. Briefly, the EV suspension with lyoprotectant was rapidly frozen, and water was removed by sublimation under vacuum conditions to preserve vesicle structure. After lyophilization, EVs were stored at −80°C, −20°C, 4°C, and room temperature(20°C) until further use. Before downstream experiments, lyophilized EVs were reconstituted in an appropriate buffer, and their concentration and quality were assessed through NTA and protein marker assessments.

### 2.11 Next-generation sequencing

Exosomal RNA was isolated using TRIzol LS Reagent (Ambion, Life Technologies, Carlsbad, CA, USA) according to the manufacturer’s instructions. RNA quality was evaluated using an Agilent 2100 Bioanalyzer or TapeStation 4000 system with an RNA 6000 Pico Chip (Agilent Technologies, Amstelveen, the Netherlands). RNA concentration was determined using either a NanoDrop 2000 spectrophotometer (Thermo Fisher Scientific) or a Qubit fluorometer (Thermo Fisher Scientific). For both control and experimental samples, small RNA libraries were prepared using the NEBNext Multiplex Small RNA Library Prep Kit (New England BioLabs, Inc., USA) following the manufacturer’s protocol. Briefly, total RNA from each sample was ligated with adaptors, and cDNA was synthesized using adaptor-specific reverse transcription primers. PCR amplification was performed, and the resulting libraries were purified using a QIAquick PCR Purification Kit (Qiagen, Hilden, Germany) and polyacrylamide gel electrophoresis. Library concentration and size distribution were assessed using a High Sensitivity DNA Assay on the Agilent 2100 Bioanalyzer (Agilent Technologies, USA). High-throughput sequencing was conducted on the Illumina NextSeq 2000 platform using single-end 75 bp reads (Illumina, San Diego, CA, USA). Raw sequencing data quality was assessed using FastQC, and adapter sequences and low-quality reads were trimmed using Trimmomatic v0.39 and Cutadapt v2.8. Processed reads were aligned to the reference genome using STAR v2.5.3a. Read quantification was performed with COMPSRA, and normalization of read counts was conducted using the TMM + CPM method implemented in the Python “conorm” package (v1.2.0).

## 3 Results

### 3.1 Phenotypic and functional characterization of the established MSCs

To confirm that the established cell line derived from hEF exhibits MSC characteristics, morphological, molecular, and functional analyses were performed (Fig. 1). Phase-contrast microscopy revealed that the cells exhibited a typical spindle-shaped, fibroblast-like morphology, consistent with the characteristic appearance of MSCs (Fig. 1A). RT-PCR analysis demonstrated that the immortalized cells maintained the expression of stemness-associated genes (SOX2, OCT4, KLF4, and MYC). Compared with Human dermal fibroblasts (HDFs) used as a negative control, the immortalized cells exhibited markedly higher expression levels of these genes, supporting the preservation of their MSC characteristics (Fig. 1B). Flow cytometric analysis showed that the cells were negative for hematopoietic lineage markers CD34, and CD45, and positive for MSC-specific markers CD73 (79.49%), CD90 (72.33%), and CD105 (68.51%) (Fig. 1C), confirming that the established cell line retains the immunophenotypic profile of MSCs. Functionally, the cells exhibited multilineage differentiation potential. Upon induction, they differentiated into adipogenic, osteogenic, and chondrogenic lineages, as evidenced by positive Oil Red O, ARS, and Alcian Blue staining, respectively (Fig. 1D). Collectively, these findings show that the established hEF-MSCs line possesses the defining morphological features, surface antigen profile, and trilineage differentiation capacity of MSCs, thereby validating its identity as an hEF-MSCs line.

**Fig. 1.**
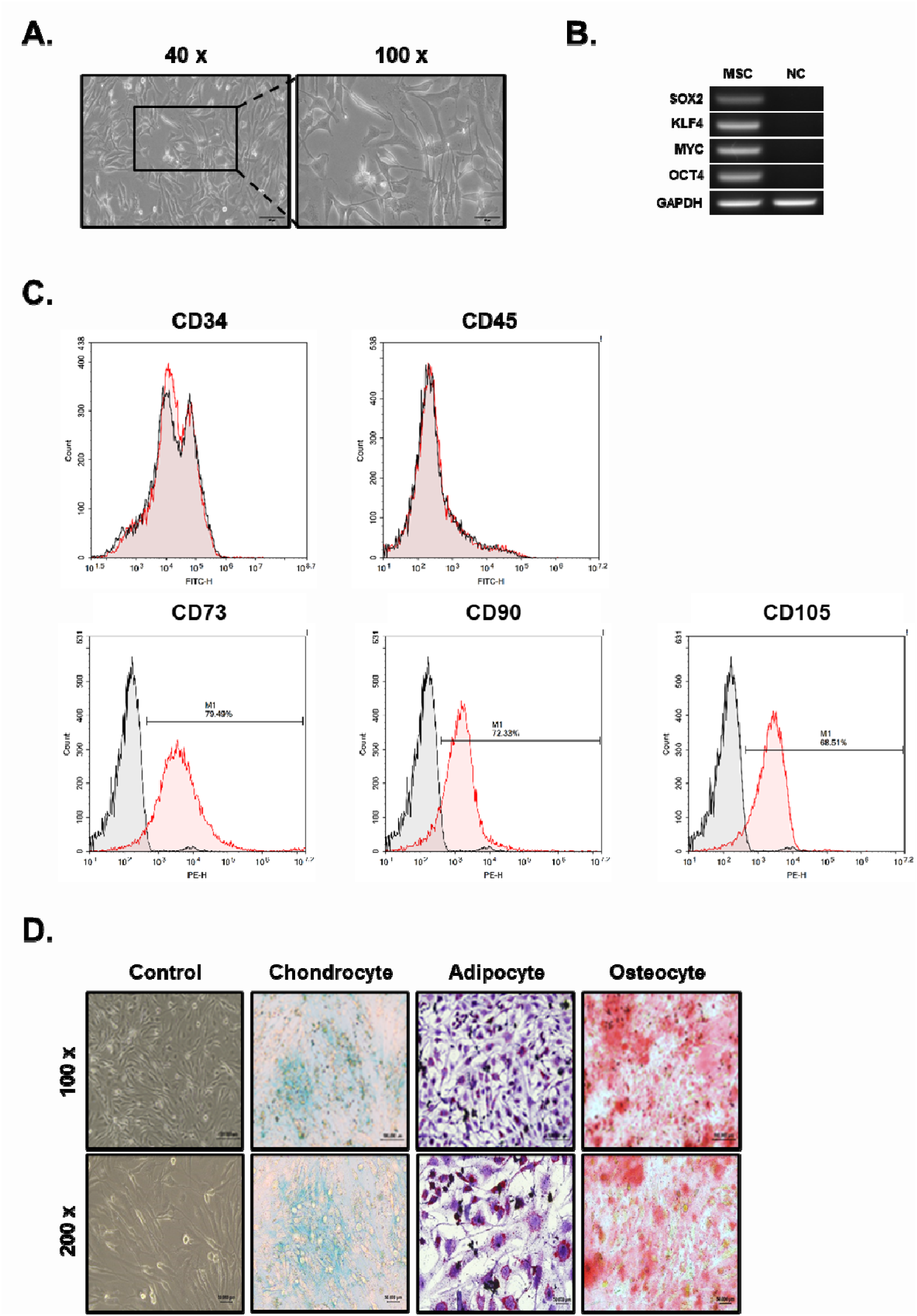
Characterization of the human epidural fat-derived mesenchymal stem cell line. (A) Representative phase-contrast images showing the typical spindle-shaped, fibroblast-like morphology of human epidural fat-derived MSCs (hEF-MSCs). (B) RT-PCR analysis demonstrating the expression of stemness-associated genes (SOX2, OCT4, KLF4, and MYC) in immortalized hEF-MSCs. Human dermal fibroblasts (HDFs) were used as a negative control (NC). (C) Flow cytometric analysis showing negative expression of CD34 and CD45. Positive expression of MSC-specific surface markers CD73 (79.49%), CD90 (72.33%), and CD105 (68.51%). (D) Multilineage differentiation potential of the MSCs confirmed through positive staining for adipogenic (Oil Red O), osteogenic (Alizarin Red S), and chondrogenic (Alcian Blue) lineages.

### 3.2 Characterization of EVs

EVs were successfully isolated from the conditioned medium of hEF-MSCs, and their defining features were confirmed through morphological and molecular analyses. TEM revealed that the isolated EVs exhibited the typical round or cup-shaped morphology of small EVs, with diameters ranging from approximately 100–200 nm (Fig. 2A). NTA further validated the vesicle size distribution and concentration, showing a mean diameter of 159.7 ± 4.7 nm and a particle concentration of approximately 1.83 × 10^11^ particles/mL (Fig. 2B). To confirm the presence of canonical EV surface markers, flow cytometric analysis was performed using antibodies against CD9, CD63, and CD81. The isolated vesicles showed strong positive signals for all three tetraspanins, with 91.74% for CD9, 91.22% for CD63, and 87.93% for CD81 (Fig. 2C). Together, these findings confirm that the isolated vesicles exhibit the characteristic morphology, size distribution, and surface marker expression of EVs derived from MSCs.

**Fig. 2.**
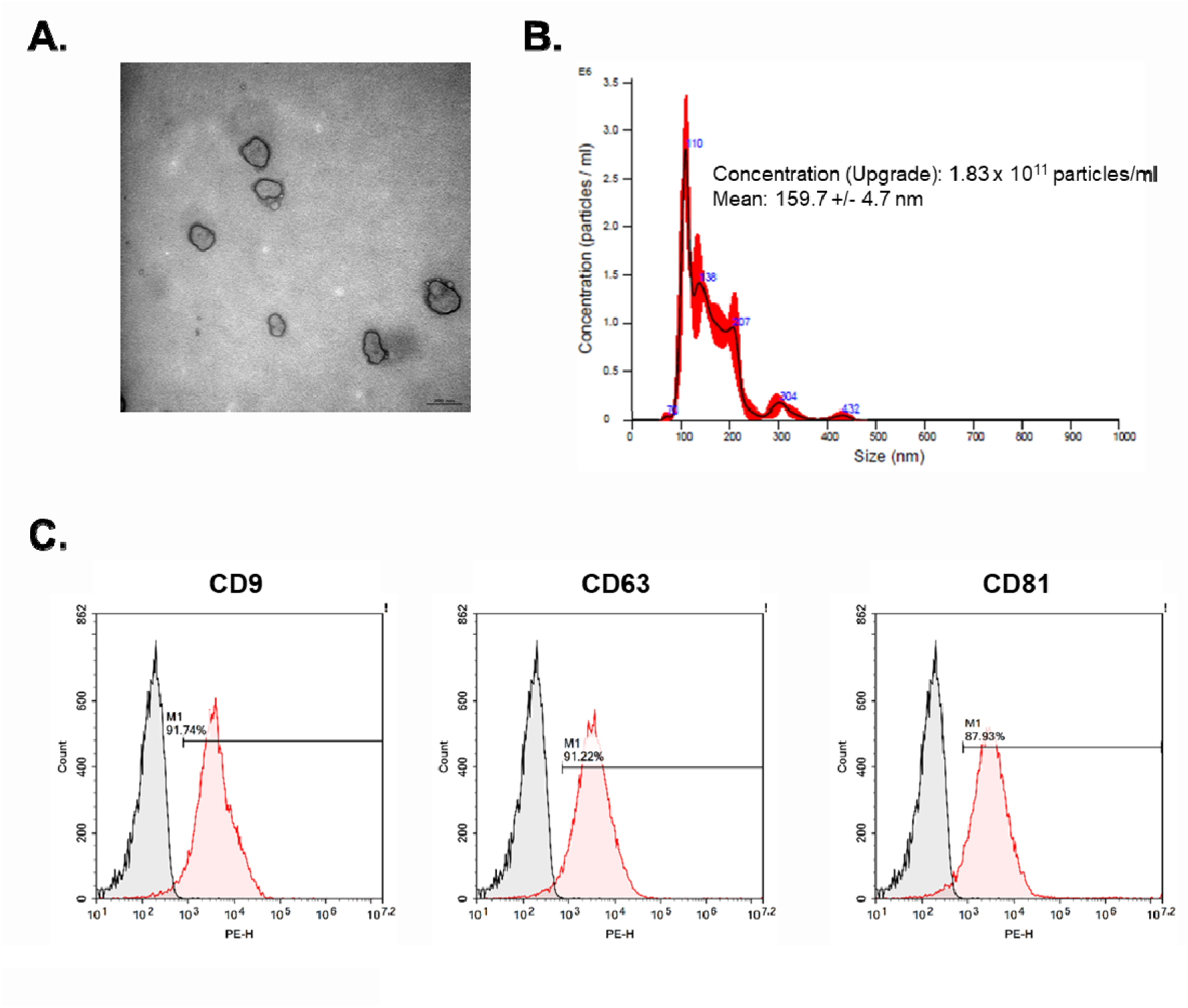
Characterization of extracellular vesicles derived from hEF-MSCs. (A) Transmission electron microscopy (TEM) images showing the characteristic cup-shaped morphology of isolated EVs. Scale bar = 200 nm. (B) Nanoparticle tracking analysis (NTA) showing a mean particle size of 159.7 ± 4.7 nm and a concentration of 1.83 × 10^11^ particles/mL. (C) Flow cytometric analysis confirming the expression of extracellular vesicle (EV)-specific surface markers CD9 (91.74%), CD63 (91.22%), and CD81 (87.93%).

### 3.3 Characterization of EVs after lyophilization

To evaluate whether lyophilization affects the characteristics of EVs derived from hEF-MSCs, lyophilized EVs were immediately reconstituted in PBS and analyzed. TEM confirmed that the reconstituted EVs retained their typical spherical or cup-shaped morphology, comparable to that of freshly isolated EVs (Fig. 3A). NTA revealed a mean particle size of 149.3 ± 4.0 nm and a concentration of approximately 1.50 × 10^11^ particles/mL, consistent with the size distribution and particle concentration of non-lyophilized EVs (Fig. 3B). To determine whether lyophilization altered canonical EV surface marker expression, flow cytometric analysis was performed. The reconstituted EVs exhibited high expression of tetraspanins CD9 (88.54%), CD63 (88.93%), and CD81 (87.21%), with no significant reduction compared to freshly isolated EVs (Fig. 3C). Overall, these results indicate that lyophilization followed by immediate PBS reconstitution preserves the morphological, physical, and molecular characteristics of EVs derived from hEF-MSCs, supporting their structural stability and suitability for downstream applications.

**Fig. 3.**
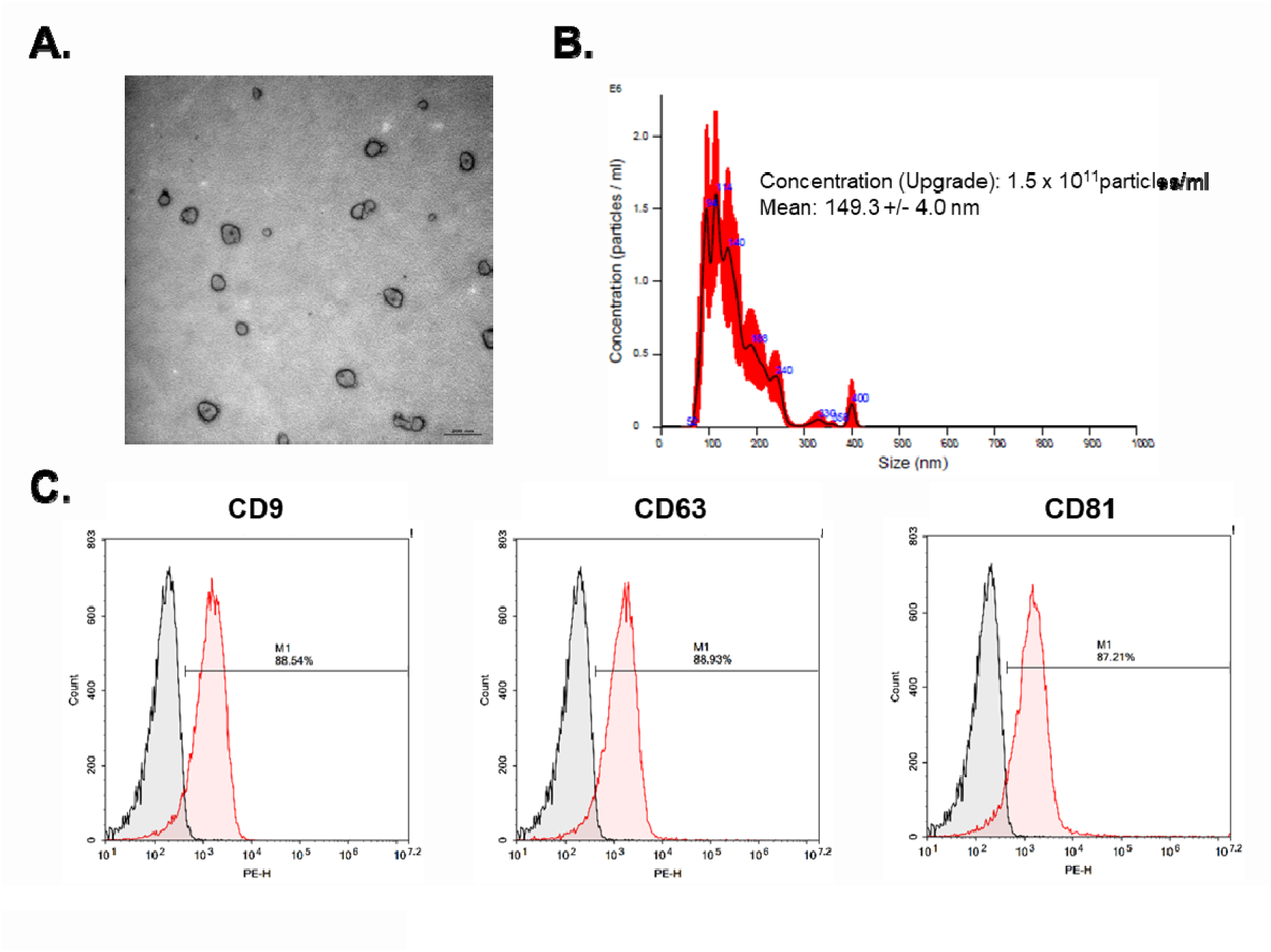
Characterization of lyophilized and PBS-reconstituted extracellular vesicles derived from hEF-MSCs. (A) TEM images showing that lyophilized and reconstituted EVs retain their characteristic spherical or cup-shaped morphology. Scale bar = 200 nm. (B) NTA showing a mean particle size of 149.3 ± 4.0 nm and a concentration of 1.50 × 10^11^ particles/mL following reconstitution. (C) Flow cytometric analysis confirming high expression of canonical EV surface markers CD9 (88.54%), CD63 (88.93%), and CD81 (87.21%), indicating that surface marker expression is preserved following lyophilization.

### 3.4 Evaluation of long-term lyophilization under storage conditions

Lyophilized EV derived from hEF-MSCs were stored at four temperatures (−80°C, −20°C, 4°C, and 20°C) for 3 months and subsequently reconstituted in PBS to evaluate their stability. Across all storage conditions, the EVs retained their fundamental physicochemical characteristics. TEM analysis revealed that EVs from all storage groups maintained their characteristic spherical or cup-shaped morphology with intact lipid bilayers, showing no evidence of structural disruption or aggregation (Fig. 4A). NTA further confirmed the preservation of particle size and concentration. The mean diameters were 179.1 ± 4.3 nm (−80°C), 145.6 ± 5.0 nm (−20°C), 158.9 ± 4.0 nm (4°C), and 127.9 ± 2.7 nm (20°C), with particle concentrations of 1.14 × 10¹¹ particles/mL (−80°C), 1.73 × 10¹¹ particles/mL (−20°C), 2.01 × 10¹¹ particles/mL (4°C), and 1.71 × 10¹¹ particles/mL (20°C) (Fig. 4B). The biochemical integrity of the EVs was assessed through flow cytometric analysis of tetraspanin markers (CD9, CD63, and CD81), all of which exhibited consistent expression across storage temperatures (Fig. 4C). Taken together, these findings indicate that lyophilized EVs derived from hEF-MSCs retain their morphological, physical, and molecular properties for at least 3 months regardless of storage temperature, supporting lyophilization as an effective strategy to overcome cold-chain limitations in EV-based therapeutic applications.

**Fig. 4.**
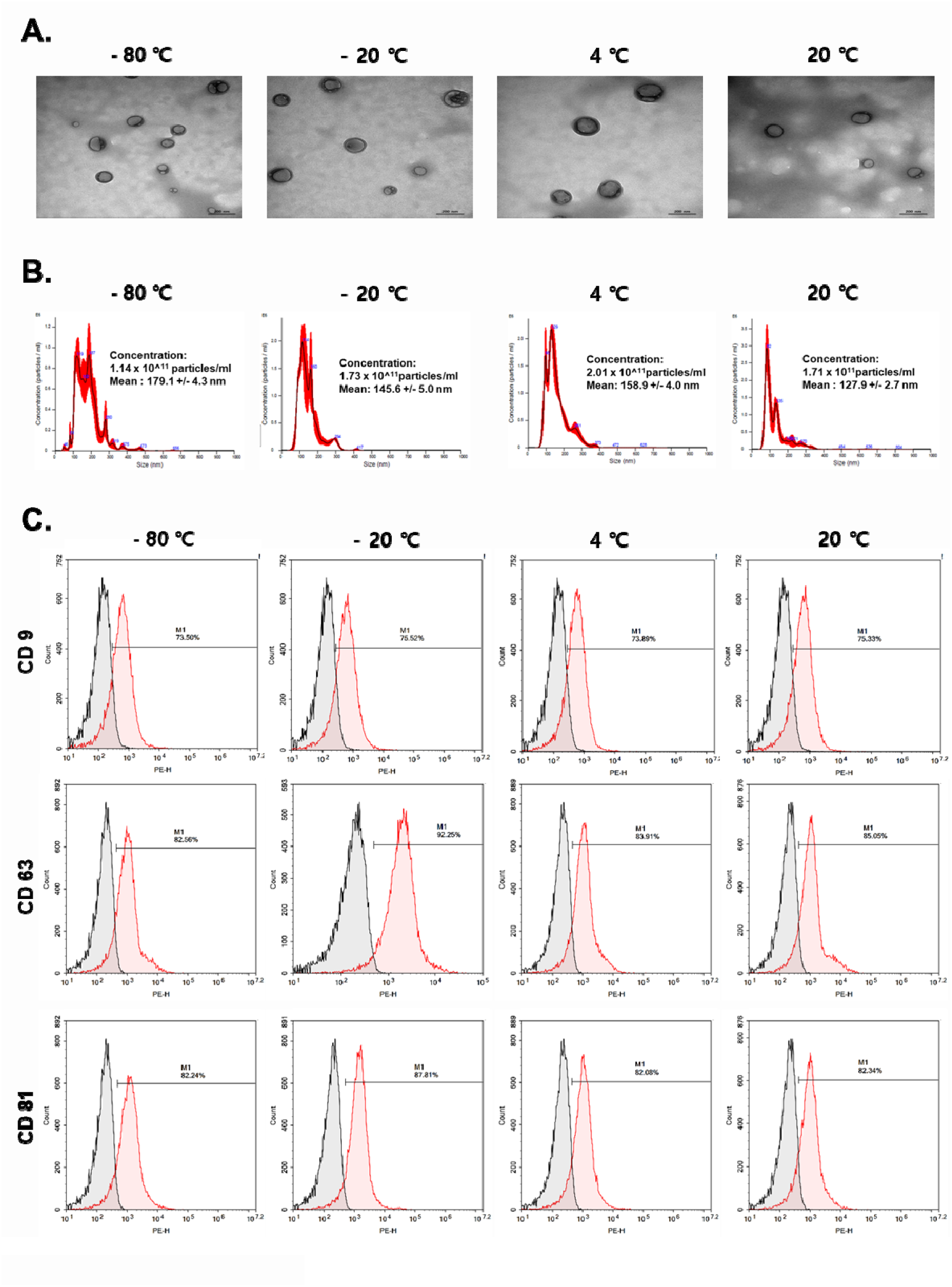
Characterization of EVs derived from hEF-MSCs. (A) Representative TEM images showing spherical morphology of EVs. Scale bars = 100 nm. (B) NTA showing particle size distribution and concentration of EVs, with mean particle size and concentration indicated for each sample. (C) Flow cytometry analysis of EV surface markers. Histogram overlays indicate positive expression of MSC-associated markers compared to isotype controls.

### 3.5 Effect of lyophilized EVs on osteogenic differentiation in MC3T3-E1 cells

The functional bioactivity of lyophilized EVs derived from hEF-MSCs and stored for 3 months at different temperatures (−80°C, −20°C, 4°C, and 20°C) was evaluated by their ability to promote osteogenic differentiation in MC3T3-E1 pre-osteoblast cells. EVs reconstituted after storage under all conditions consistently enhanced osteogenic differentiation compared with untreated controls, indicating that their activity was preserved regardless of storage temperature. Cell viability following EV treatment was assessed using the MTT assay (Fig. 5A). MC3T3-E1 cells were treated with 1 × 10^8^ EV particles. No significant cytotoxicity was observed in any EV-treated group compared with the control, confirming that the observed increases in osteogenic differentiation were not attributable to cell death. Early osteoblast marker gene expression, including *DLX5* and *RUNX2*, was assessed using RT-PCR (Fig. 5B). Gel electrophoresis showed distinct upregulation of both genes in all EV-treated groups, regardless of storage temperature (−80°C, −20°C, 4°C, and 20°C). These findings indicate that neither lyophilization nor storage conditions impaired the intrinsic pro-osteogenic signaling capacity of the EVs. Functional osteogenic activity was further validated through ALP staining and ARS staining. ALP staining (Fig. 5C) revealed markedly increased ALP activity in all EV-treated groups relative to the control, indicating effective induction of early osteogenic differentiation and preparation for matrix mineralization. ARS staining (Fig. 5D) showed robust and widespread calcium deposition in EV-treated cells compared with the control, confirming that the EVs promoted late-stage osteogenic maturation and mineralization regardless of storage temperature. Collectively, these results show that lyophilized EVs derived from hEF-MSCs retain their full osteogenic activity—including upregulation of *DLX5* and *RUNX2*, enhanced ALP activity, and promotion of calcium deposition—after 3 months of storage across a wide temperature range (−80°C to 20°C), without inducing cytotoxicity. These findings establish lyophilization as an effective strategy for the long-term preservation and transport of EVs at room temperature(20°C) for therapeutic applications.

**Fig. 5.**
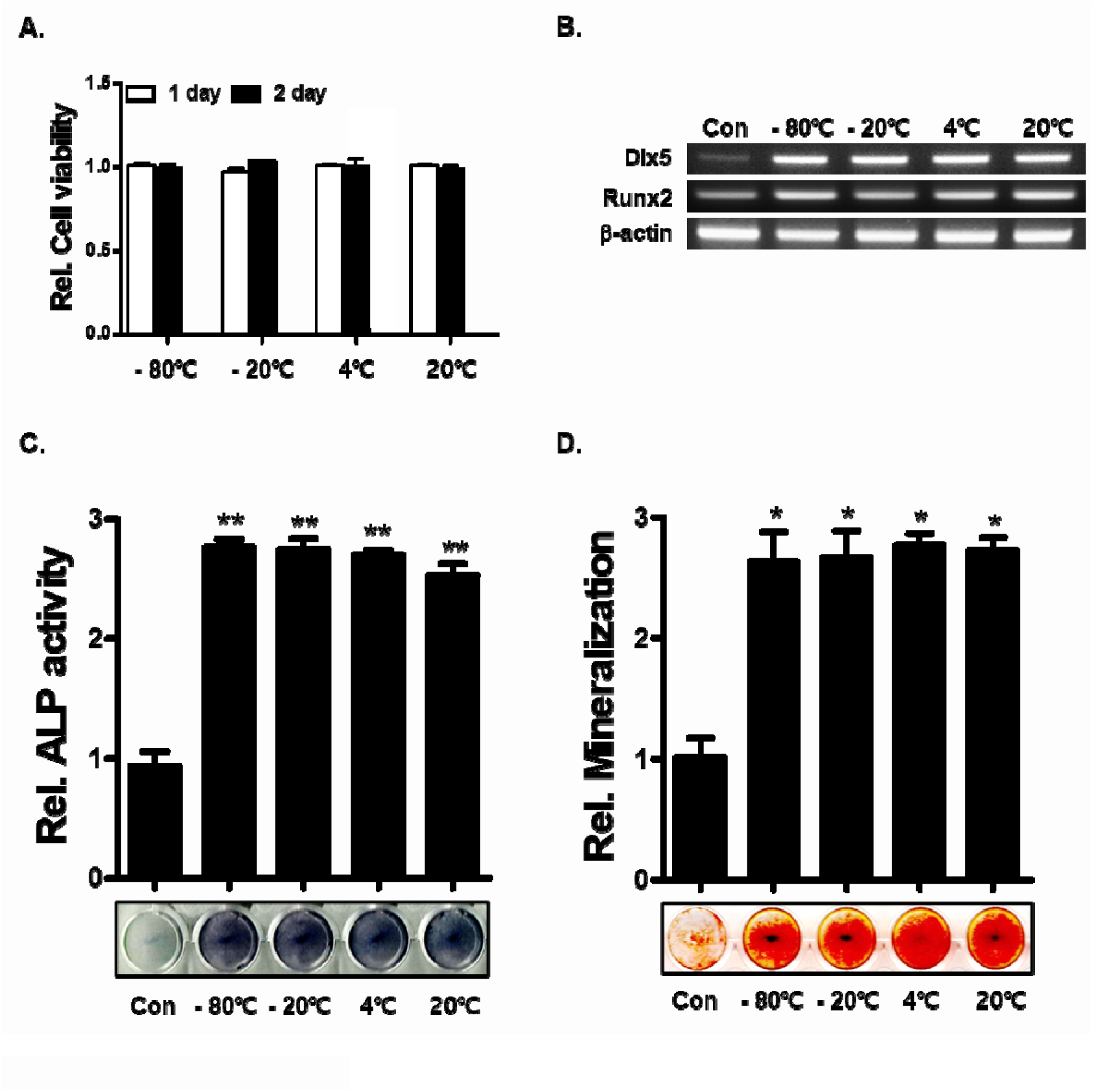
EF-derived EVs promote proliferation and osteogenic differentiation of MC3T3-E1 cells. (A) MC3T3-E1 cells were treated with epidural fat-derived EVs at 1 × 108 particles/mL for 1 and 2 days. Cell viability was assessed using the MTT assay, indicating no significant cytotoxicity. (B) Osteogenic gene expression of *DlX5* and *RUNX2* was assessed through RT-PCR following 2 days of EV treatment, showing upregulation of early osteogenic markers. (C, D) Alkaline phosphatase at 7 days and Alizarin Red S staining at 21 days following osteogenic induction with EV treatment revealed enhanced early differentiation and matrix mineralization in EV-treated cells compared with controls. Statistical significance was determined using the Student’s *t*-test (\**p* < 0.05, \*\**p* < 0.01).

### 3.6 Analysis of miRNA in EVs after long-term lyophilization

To evaluate the stability of the internal miRNA cargo of lyophilized EVs derived from hEF-MSCs, gene expression profiling was performed using next-generation sequencing (NGS) after 3 months of storage at −80°C, −20°C, 4°C, and 20°C. All storage groups were compared with EVs stored at room temperature(20°C). miRNA expression patterns were first examined using scatter plots and volcano plots to compare transcriptomic profiles across storage groups. Scatter plots showed that most miRNAs aligned tightly along the central diagonal (y = x), indicating a high degree of similarity among the groups. The absence of substantial deviations from the diagonal suggests that the miRNA composition of the EVs remained largely unchanged despite prolonged storage across different temperatures (Fig. 6A). Volcano plots further illustrated the magnitude (log fold change) and statistical significance (−log *p*-value) of differences in miRNA abundance among groups. Across all comparisons, only a small subset of miRNAs exhibited statistically significant changes, while the majority remained within the nonsignificant central region. These results indicate that storage temperature did not induce significant changes in EV miRNA cargo (Fig. 6B). Global similarity in miRNA profiles was assessed using principal component analysis (PCA) (Fig. 6C). The 3D PCA plot—based on PC1, PC2, and PC3—showed that samples from all storage temperatures clustered closely together, with minimal dispersion. Replicate samples within each group exhibited very low variability, indicating excellent reproducibility. Notably, the spatial proximity of clusters across all temperature conditions indicates that EVs stored at 20°C retained miRNA expression profiles that were statistically indistinguishable from those stored at −80°C. Collectively, these findings indicate that lyophilized EV derived from hEF-MSCs maintain highly stable miRNA cargo profiles after 3 months of storage across temperatures ranging from −80°C to room temperature(20°C). These results align with the observed preservation of the physical characteristics and functional bioactivity of EVs, providing strong molecular evidence that lyophilization is an effective approach for long-term EV preservation and room-temperature distribution.

**Fig. 6.**
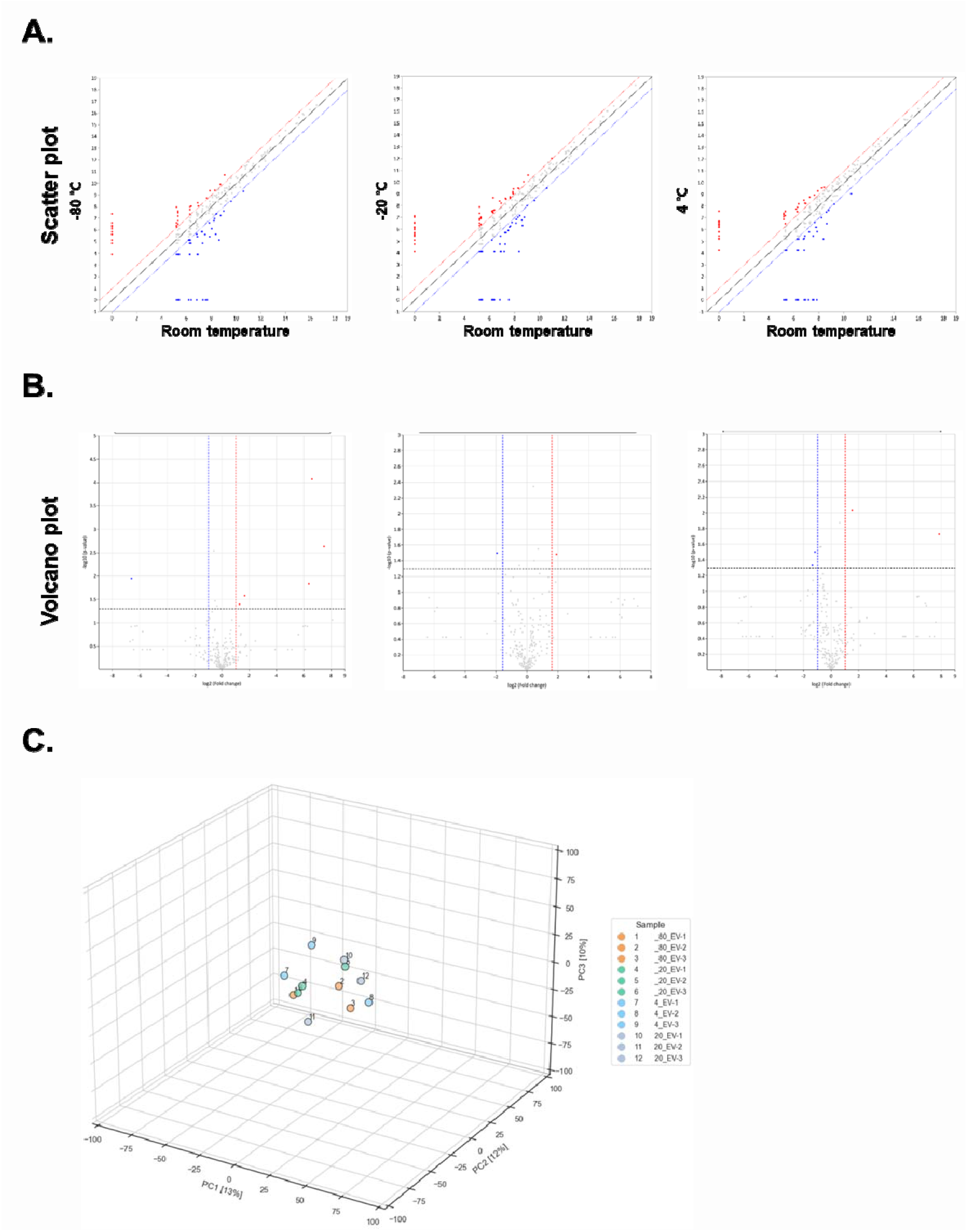
Transcriptomic analysis of lyophilized EVs derived from hEF-MSCs following 3 months of storage at different temperatures. (A) Scatter plot comparing miRNA expression profiles between room-temperature EVs and lyophilized EVs stored at −80°C, −20°C, and 4°C. (B) Volcano plots showing differential miRNA expression in lyophilized EVs after 3 months of storage. Horizontal dashed lines represent statistical significance thresholds (−log_10_ *p*-value), and vertical dashed lines indicate fold-change thresholds. (C) Three-dimensional principal component analysis of next-generation sequencing data showing tight clustering of biological replicates within each storage condition.

## 4 Discussion

EVs derived from MSCs have emerged as potent modulators of intercellular communication with significant therapeutic potential, particularly in regenerative medicine (Fuloria et al., 2021; Matsuzaka and Yashiro, 2022; Thalakiriyawa et al., 2022). Among various applications, their ability to enhance bone regeneration and support osteogenic differentiation has garnered substantial interest due to the need for effective treatments for osteoporosis, delayed fracture healing, and other skeletal disorders (Al-Sharabi et al., 2024; Zhang et al., 2020). Despite these therapeutic prospects, the clinical translation of MSC-derived EVs has been constrained by the challenges associated with long-term storage, stability, and the preservation of functional activity. Conventional cryopreservation at −80°C or 4°C helps maintain EV structural integrity but imposes practical challenges, including high operational costs, reliance on continuous cold-chain infrastructure, and limited feasibility for large-scale global distribution (Ahmadian et al., 2024; Gelibter et al., 2022; Klymiuk et al., 2024; Lőrincz Á et al., 2014; Yuan et al., 2021). In this study, we evaluated lyophilization as an alternative storage strategy to address these limitations. Using EVs derived from immortalized hEF-MSCs, we assessed their physicochemical characteristics, biological activity, and molecular cargo stability following long-term storage across multiple temperature conditions. The immortalized hEF-MSCs retained the expression of mesenchymal stem cell-associated markers, indicating preservation of their MSC phenotype, consistent with previous reports demonstrating that immortalization does not necessarily compromise MSC characteristics (Lee et al., 2019). The maintenance of these MSC characteristics supports the use of immortalized hEF-MSCs as a dependable EV source and strengthens the translational relevance of the present findings.

Our findings show that lyophilized EVs retain morphological, molecular, and functional characteristics comparable to those of freshly isolated EVs. TEM and NTA indicated that the EVs maintained their canonical spherical or cup-shaped morphology, with no evidence of structural disruption or aggregation following lyophilization and reconstitution. Particle size and concentration remained largely preserved, suggesting that the lyophilization process, when combined with appropriate cryoprotectants such as 8.5% sucrose and 20 mM HEPES, effectively mitigates vesicle fusion, collapse, or aggregation during the drying process. These results align with previous studies, which showed that lyoprotectants—particularly sugars such as sucrose, trehalose, and mannitol—help stabilize the lipid bilayer and preserve the vesicular architecture by forming a protective glassy matrix that reduces mechanical stress during sublimation (Ahmadian et al., 2024; Boafo et al., 2022; Sikandar et al., 2024). In this study, sucrose provided structural stabilization while also offering benefits associated with its biocompatibility, FDA approval, and suitability for GMP-compliant manufacturing, thereby supporting its translational feasibility for clinical applications (Castañeda Ruiz et al., 2022; Ionova and Wilson, 2020; Lo Presti et al., 2024; Sarvepalli et al., 2025).

Flow cytometric analysis showed that lyophilization did not alter the expression of canonical EV surface markers, including CD9, CD63, and CD81. Notably, the EVs retained high levels of tetraspanin positivity after lyophilization, highlighting the preservation of membrane protein composition, which is critical for recognition and uptake by recipient cells. These findings were consistent across storage temperatures ranging from −80°C to room temperature (20°C) over a 3-month period, suggesting that lyophilized EVs can remain stable without reliance on the stringent cold-chain conditions typically required for EV-based therapies. The maintenance of these physical and molecular properties is crucial, as alterations in vesicle morphology or marker expression may affect biodistribution, cellular uptake, and downstream biological activity.

Beyond maintaining structural integrity, the functional bioactivity of lyophilized EVs was rigorously evaluated using MC3T3-E1 pre-osteoblast cells. Across all storage conditions, reconstituted EVs consistently promoted osteogenic differentiation, as evidenced by the upregulation of key osteogenic-related genes *DLX5* and *RUNX2*, enhanced ALP activity, and extensive calcium deposition detected with ARS staining. These findings indicate that lyophilization preserves both the physical properties and the intrinsic biological activity of MSC-derived EVs, allowing them to retain their capacity to influence recipient cell behavior. Furthermore, MTT assays confirmed that EV treatment did not induce cytotoxicity under any storage conditions, confirming that the observed enhancement in osteogenic differentiation was attributable to preserved EV bioactivity rather than nonspecific cellular stress or reduced cell viability.

Our study also shows that the molecular cargo of EVs, particularly their miRNAs, remains highly stable following long-term storage. NGS analysis revealed that the miRNA profiles of lyophilized EVs remained highly consistent across all storage temperatures, including room temperature (20°C). This conclusion was supported by minimal deviations in scatter and volcano plots and by tight clustering in PCA. This is of significant translational importance because the miRNA content of EVs mediates many of their paracrine effects, including the regulation of osteogenic signaling pathways. The retention of a stable miRNA repertoire suggests that lyophilization effectively prevents the degradation of nucleic acid cargo, reinforcing the suitability of this approach for both clinical and research applications. Collectively, the sustained preservation of EV morphology, surface-marker expression, functional activity, and miRNA profiles underscores the robustness of lyophilization as a comprehensive EV storage approach.

The ability to store EVs at multiple temperatures without compromising their functional properties represents a substantial advancement in regenerative medicine. Lyophilization offers a practical and scalable solution for the storage and distribution of EV-based therapeutics by mitigating reliance on cold-chain logistics and reducing associated operational costs. This is particularly relevant for widespread clinical implementation, where storage at conventional cryogenic temperatures is not always feasible, especially in resource-limited settings. Furthermore, the preservation of EV bioactivity across temperature conditions may facilitate its incorporation into off-the-shelf therapeutic products, enabling clinical use without the need for specialized storage infrastructure.

Previous studies have reported potential limitations in the stability of EVs under lyophilization, including inconsistent preservation of vesicle morphology and functional activity depending on the selected lyoprotectant and storage conditions (Frank et al., 2018; Lyu et al., 2022; Susa et al., 2023; Trenkenschuh et al., 2022). Our study addresses these gaps by systematically evaluating a clinically relevant EV source hEF-MSCs using an optimized sucrose-based cryoprotectant system and assessing both short-term structural features and long-term functional outcomes. By integrating detailed morphological, molecular, and functional analyses, including comprehensive NGS-based profiling of miRNA cargo, we provide robust evidence that supports the generalizability of lyophilized EVs derived from hEF-MSCs as a stable therapeutic approach.

Beyond its implications for EV-based regenerative therapies, this study advances the broader field of nanomedicine by offering insights into the stabilization of biologically derived vesicles for clinical applications. Unlike synthetic lipid nanoparticles, which often require complex modifications such as PEGylation or targeting ligands to achieve stability and controlled biodistribution, natural EVs offer inherent biocompatibility, efficient cellular uptake, and the capacity to transport diverse bioactive molecules (Bader et al., 2024; Jeyaram and Jay, 2017; Tenchov et al., 2023; Villa et al., 2019). Lyophilization further enhances the clinical feasibility of these natural advantages by enabling long-term storage and transport without compromising functionality, thus addressing a key barrier to the translational application of EV-based therapeutics.

## 5 Conclusions

Our study indicates that lyophilization of EV derived from hEF-MSCs using sucrose as a cryoprotectant preserves their morphological, molecular, and functional integrity for at least 3 months across a broad range of storage temperatures. These lyophilized EVs retain their capacity to promote osteogenic differentiation without inducing cytotoxicity and maintain a highly stable miRNA cargo, highlighting the potential of lyophilization as a practical, scalable preservation strategy. These findings provide strong evidence supporting the feasibility of off-the-shelf EV therapeutics for bone regeneration and establish a foundation for further clinical translation and large-scale production of MSC-derived EVs. Future studies should explore longer storage durations, evaluate *in vivo* biodistribution and therapeutic efficacy, and assess the applicability of this preservation approach to EVs derived from diverse cellular sources. Collectively, this study highlights the transformative potential of lyophilized EVs as stable, effective, and clinically deployable agents in regenerative medicine.

## Ethics Declarations Consent to participate

This study was approved by the Institutional Review Board of Yeungnam University Medical Center (IRB No. 2017-07-032).

## Consent to Publish

Not applicable.

## Clinical trial number

Not applicable.

## Availability of data and material

The data that support the findings of this study are available from the corresponding author upon reasonable request.

## Author contributions

Young-Ju Lim and Wook Tae Park contributed equally to this work. Young-Ju Lim, Wook Tae Park, and Gun Woo Lee were involved in the conception and design of the study. Young-Ju Lim, Wook Tae Park, Joo-Hee Choi performed the experiments and analyzed and interpreted the data. Sangbum Park, Min-Soo Seo and Gun Woo Lee provided critical supervision and intellectual input. Young-Ju Lim and Wook Tae Park drafted the manuscript. All authors contributed to revising the manuscript critically for important intellectual content, approved the final version to be published, and agree to be accountable for all aspects of the work.

